# “*Citrobacter bitternis*” Ko et al., 2015 and “*Kluyvera intestini*” Tetz and Tetz, 2016 are heterotypic synonyms of *Phytobacter diazotrophicus* Zhang et al., 2017

**DOI:** 10.1101/2025.08.11.669625

**Authors:** Fabio Rezzonico, Theo H. M. Smits

**Affiliations:** Environmental Genomics and Systems Biology Research Group, Institute of Natural Resource Sciences (IUNR), Zurich University of Applied Sciences ZHAW, Wädenswil, Switzerland

**Keywords:** *Enterobacteriaceae*, misidentification, taxonomic revision, comparative genomics

## Abstract

Members of *Phytobacter*, a recently described genus within the family Enterobacteriaceae, are still frequently misclassified as species of other genera. Here, we present genomic evidence that the taxa *“Citrobacter bitternis”* and *“Kluyvera intestini”*, as cited in the literature and listed in NCBI databases, are heterotypic synonyms of *Phytobacter diazotrophicus*. Comparative analyses of 16S rRNA gene phylogeny, core genome phylogeny, average nucleotide identity (ANI), and digital DNA–DNA hybridization (dDDH) consistently place the type strains of both *“C. bitternis”* and *“K. intestini”* within the species *P. diazotrophicus*. Recognizing these taxa as synonyms will help resolve persistent taxonomic inconsistencies and reduce misidentification in clinical and environmental microbiology.

## Introduction

Taxonomic changes within the family *Enterobacteriaceae* are still ongoing, mostly driven by the continuous generation of new genome sequences [1, 2]. The genus *Phytobacter* was first described in 2008 [3], while the name was validated only in 2017. Due to its novelty, the genus is still prone to have pre-dating identification attempts under different names [4]. This confusion, albeit partially resolved in recent years [4], still persists.

In recent years, the fundamental knowledge about the genus *Phytobacter* and its species [5, 6] has been expanded, thereby improving the identification procedures of their members using qPCR [7] and MALDI-TOF MS [8]. This effort has led to the acceptance of the species as a rare, but emerging human pathogen that can carry clinically important antimicrobial resistance genes [9, 10]. Recently characterized isolates belonging to the genus are usually correctly identified, especially when having a complete genome sequence [11-13]. However, there are still incidences in which the gathered knowledge is ignored or misinterpreted.

While working on comparative genomics of the genus *Phytobacter*, the presence of entries with a different name having no taxonomic standing within the same species can not only be irritating, but also be the source of further misidentification issues if these are used as references. Two cases are reported below.

### The case “*Kluyvera intestini*”

The name “*Kluyvera intestini*” was only used in two genome announcements [14, 15], mainly describing the genome sequence of the specific isolate GT-16. Already in 2019 [16], it was reported that “*K. intestini*” GT-16 rather clustered with genomes of *Phytobacter diazotrophicus*, and this was confirmed in a later study on taxonomic confusion [4]. Nevertheless, this genome is still listed in National Center for Biotechnology Information (NCBI) GenBank under the incorrect name, while maintaining it as a “not valid description” in the List of Prokaryotic names with Standing in Nomenclature (LPSN). In 2017, the genome was used to support the transfer of “*Enterobacter massiliensis*” to the novel genus “*Metakosakonia*” [2], which later was then unified with the genus *Phytobacter* [17].

### The case “Citrobacter bitternis”

The species “*Citrobacter bitternis*” [18] was initially described relying solely on 16S rRNA gene sequences, which were exclusively compared with sequences of *Citrobacter* spp. This is not sufficient for the delineation of a new species within the *Enterobacteriaceae*, as the 16S rRNA gene is too conserved to serve as a sole biomarker for this family [19]. Multi-locus sequencing analysis (MLSA), as implemented for species delineation since 2008 for different *Enterobacterales* [20, 21], would have resulted in a more accurate identification.

The recent addition of the genome of type strain “*Citrobacter bitternis*” JCM 30009 (accession no. JBHSRG01) has issued the NCBI to create another level of confusion. New genomes, correctly identified as *P. diazotrophicus* by the submitters, are since then marked as “unverified source organism” due to the comparison to the “*C. bitternis*” genome. However, according to the LPSN, the name “*C. bitternis*” is not validly described as well.

### Evidence for an erroneous assignment of “*C. bitternis*” and “*K. intestini*”

The genomes of both “*C. bitternis*” JCM 30009 and “*K. intestini*” GT-16 were investigated for different taxonomic parameters. Currently, 16S rRNA gene sequences of six different strains belonging to the species “*C. bitternis*” are available in GenBank. Additionally, the consensus 16S rRNA gene sequences for “*C. bitternis*” JCM 30009 and “*K. intestini*” GT-16 were extracted from the respective genomes. In a phylogenetic tree with the 16S rRNA gene of type strains of different *Enterobacteriaceae*, all mentioned “*C. bitternis*” 16S rRNA gene sequences cluster together with the 16S rRNA gene sequence of *P. diazotrophicus*, including its type strain DSM 17806^T^ (**Figure 1**).

**Figure 1:**
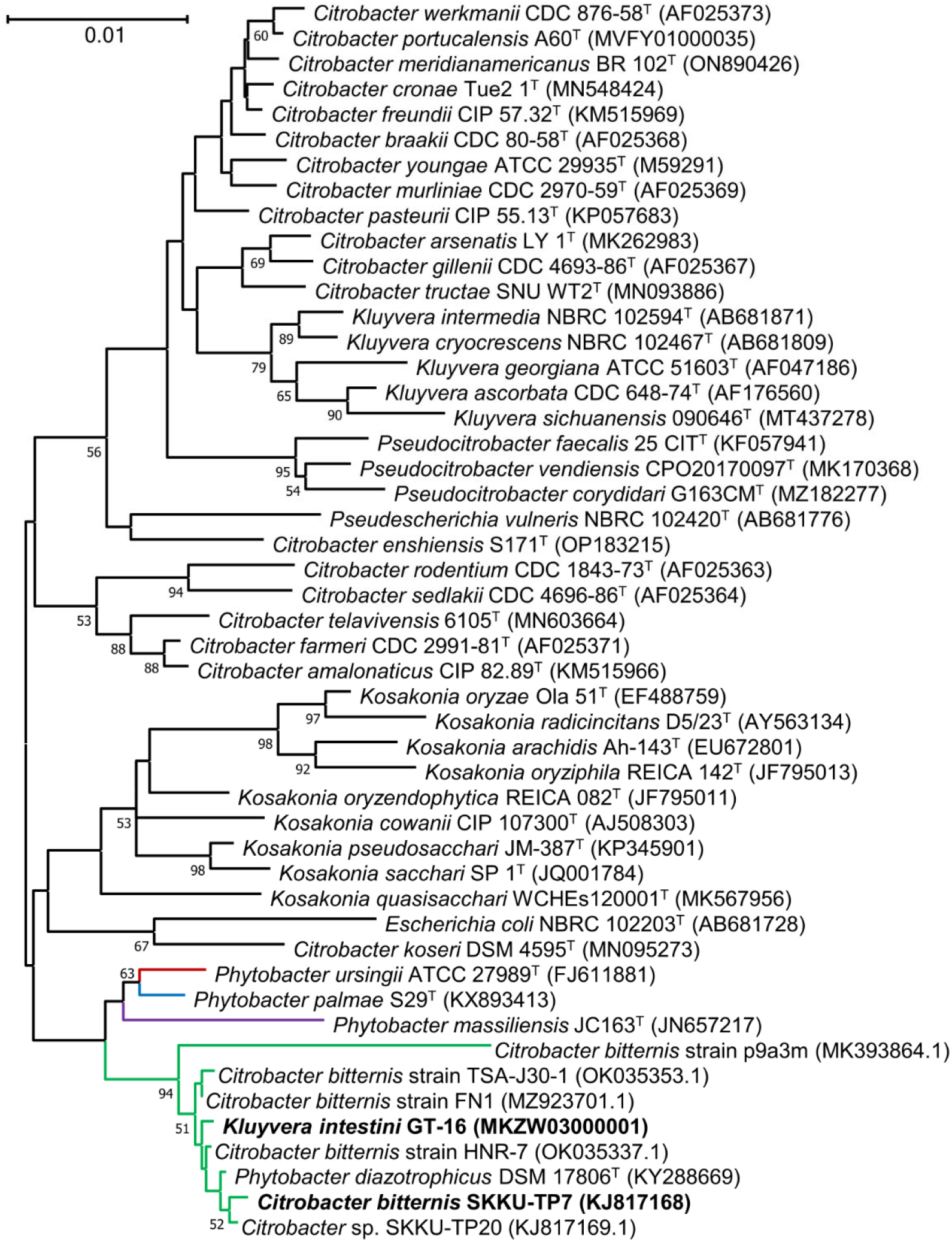
Phylogenetic tree of 16S rRNA genes inferred using the Neighbor-Joining method. The percentage of replicate trees in which the associated taxa clustered together in the bootstrap test (1000 replicates) are shown below the branches. Only percentages above 50% are considered. The GenBank accession number for the 16S rRNA gene of each strain is indicated in brackets. Lineages of *Phytobacter* spp. are indicated in color according to Smits et al., 2022 [4]. The positions of the strains “*C. bitternis*” JCM 30009 and “*K. intestini*” GT-16 are indicated in bold. test dataset.

It was noted in a recent publication [22], in which a *Phytobacter palmae* strain was identified, that in the Type (Strain) Genome Server (TYGS) [23], the genome sequences of both “*C. bitternis*” and “*K. intestini*” are clustering closely with the genomes of *P. diazotrophicus* TA9759 [11] and type strain *P. diazotrophicus* DSM 17806^T^ [5]. Accordingly, in a core genome tree generated with EDGAR [24] that included the type strains of species from *Kluyvera, Citrobacter, Phytobacter* and some other genera, the genomes of “*C. bitternis*” JCM 30009 and “*K. intestini*” GT-16 do not cluster with the corresponding type strains of *Kluyvera* or *Citrobacter* but are rather included within *P. diazotrophicus* (**Figure 2**).

**Figure 2:**
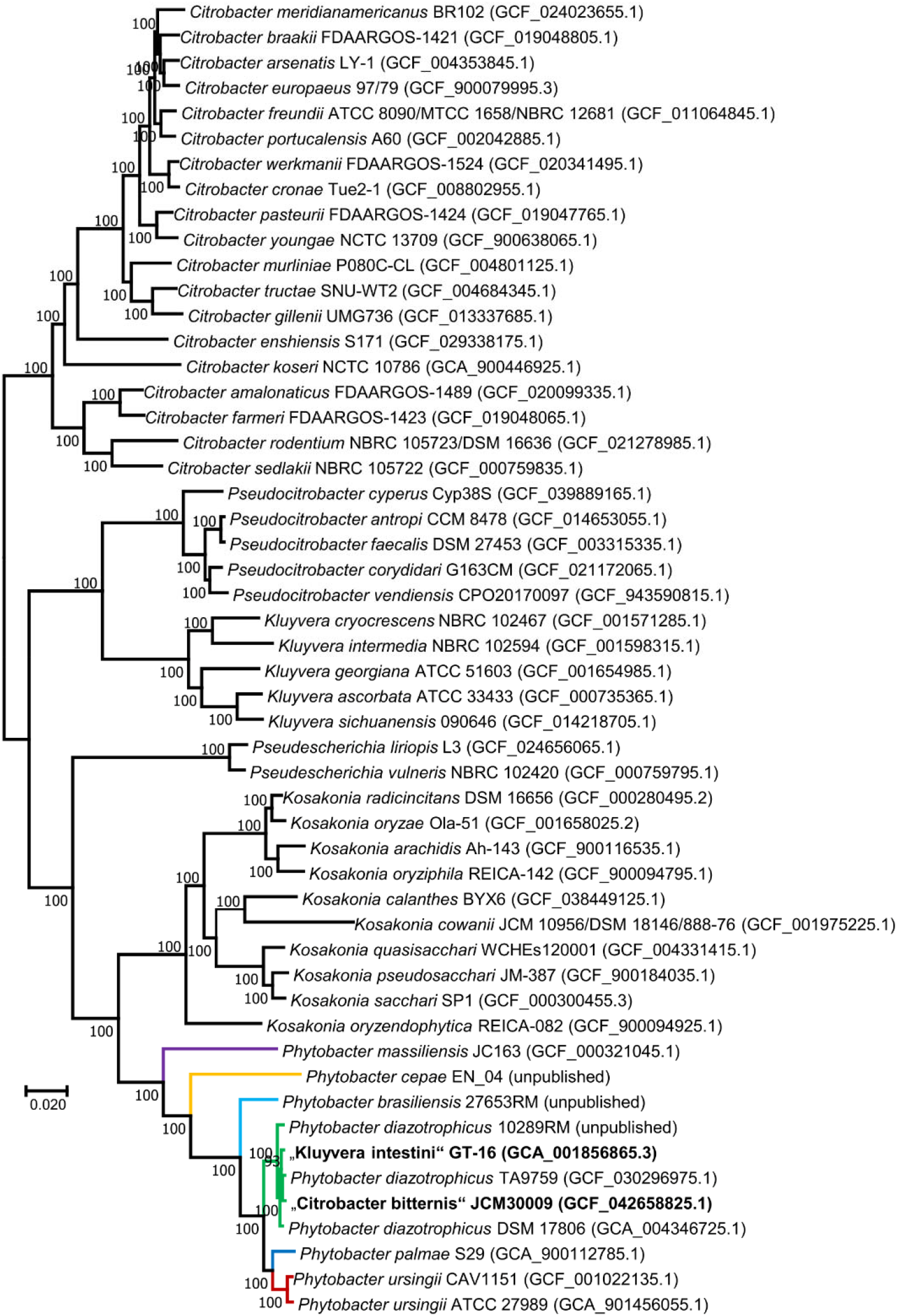
Core genome tree of type strain genomes of selected *Enterobacteriaceae*, including the genera *Phytobacter, Citrobacter* and *Kluyvera*, as created by EDGAR [24]. The assembly number of each respective genome is indicated in brackets. Values indicated at nodes are local support values computed by FastTree using the Shimodaira-Hasegawa test. Lineages of *Phytobacter* spp. are indicated in color according to Smits et al., 2022 [4]. The positions of the genomes of strains “*C. bitternis*” JCM 30009 and “*K. intestini*” GT-16 are indicated in bold.

Determination of the overall genome relatedness indices (OGRIs) confirmed that the average nucleotide identities (ANI) determined with FastANI in EDGAR [24, 25] and digital DNA:DNA hybridization values (using GGDC) [26, 27] for the genomes of both “*K. intestini*” GT-16 and “*C. bitternis*” JCM 30009 are within the range of species delineation in *P. diazotrophicus* DSM 17806^T^ (**Figure 3**). Therefore, the isolates “*C. bitternis*” JCM 30009 and “*K. intestini*” GT-16 should thus rather be considered as isolates of *P. diazotrophicus*.

**Figure 3:**
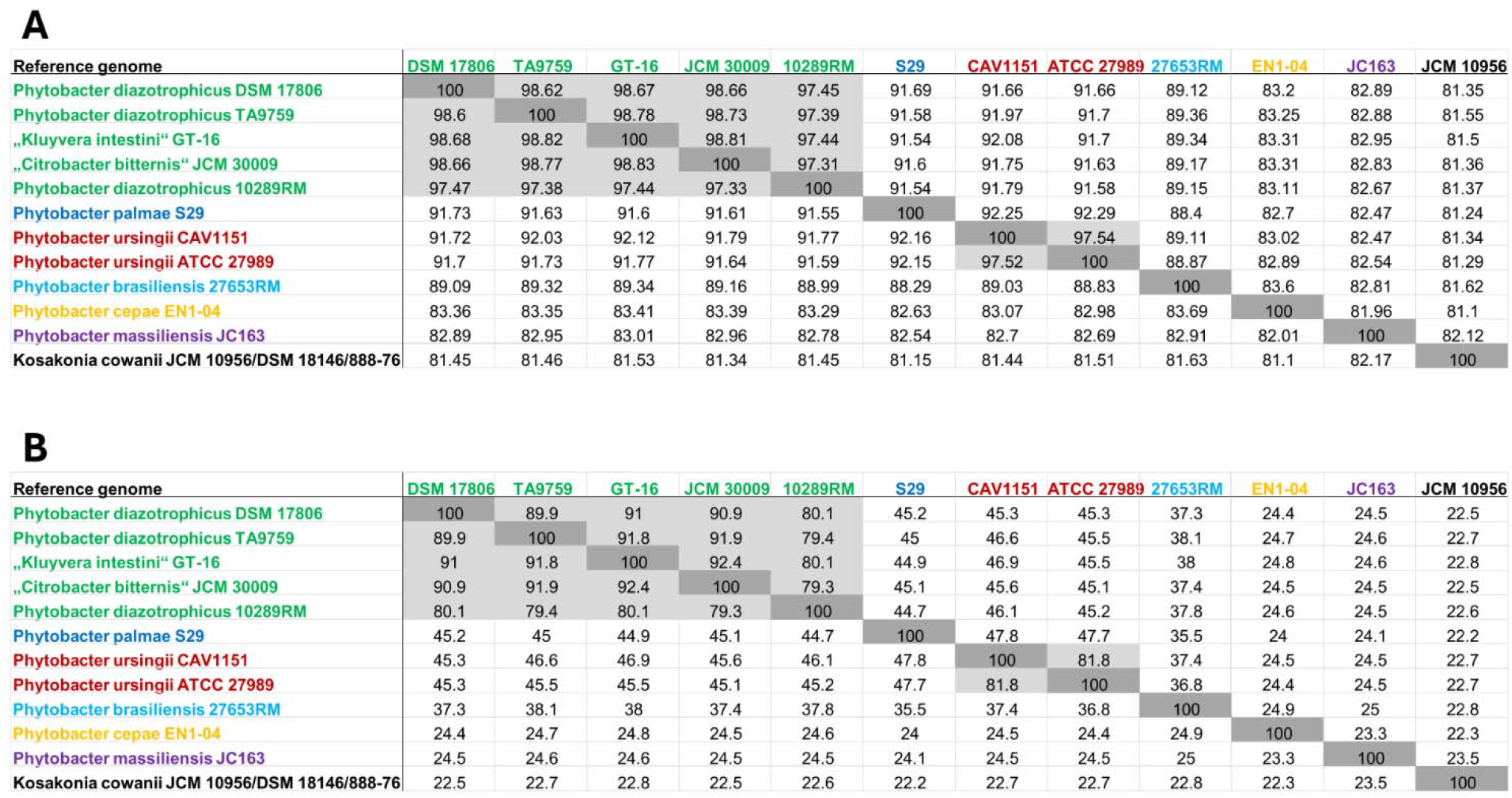
Average nucleotide identity (A) and Genome-to Genome DNA-DNA Hybridization (B) values for all *Phytobacter* spp. and outgroup *Kosakonia cowanii* JCM 10956^T^. Lineages of *Phytobacter* spp. are indicated in color according to Smits et al., 2022 [4].

## Conclusion

Based on the evidence presented in this manuscript, we propose that “*K. intestini*” Tetz and Tetz, 2016 and “*C. bitternis*” Ko et al., 2015 should be considered as heterotypic synonyms of *P. diazotrophicus*.

